# Caloric restriction mitigates age-associated hippocampal differential CG and non-CG methylation

**DOI:** 10.1101/175810

**Authors:** Niran Hadad, Archana Unnikrishnan, Jordan A. Jackson, Dustin R. Masser, Laura Otalora, David R. Stanford, Arlan Richardson, Willard M. Freeman

## Abstract

Brain aging is marked by cognitive decline and susceptibility to neurodegeneration. Caloric-restriction (CR) increases neurogenesis, improves memory function, and protects from age-associated neurological disorders. Epigenetic mechanisms, including DNA methylation, are vital to normal CNS cellular and memory functions, and are dysregulated with aging. The beneficial effects of CR have been proposed to work through epigenetic processes, but this is largely unexplored. We therefore tested whether life-long CR prevents age-related DNA methylation changes in the brain. Hippocampal DNA from young (3 months) and old (24 months) male mice fed *ad libitum* and 24 month old mice fed a 40% calorie-restricted diet from 3 months of age were examined by genome-wide bisulfite sequencing to measure methylation with base-specificity. Over 27 million CG and CH (non-CG) sites were examined. Of the ~40,000 differentially methylated CGs (dmCGs) and ~80,000 CHs (dmCHs) with aging, >1/3 were prevented by CR and were found across genomic regulatory regions and gene pathways. CR also caused alterations to CG and CH methylation at sites not differentially methylated with aging, and these CR-specific changes demonstrated a different pattern of regulatory element and gene pathway enrichment than those affected by aging. CR-specific DNMT1 and TET3 promoter hypermethylation corresponded to reduced gene expression. These findings demonstrate that CR attenuates age-related CG and CH hippocampal methylation changes, in combination with CR-specific methylation that may also contribute to the neuroprotective effects of CR. The prevention of age-related methylation alterations is also consistent with the pro-longevity effects of CR working through an epigenetic mechanism.

## 1. Introduction

With increased life-expectancy, incidence of age-related neurodegenerative diseases such as Alzheimer’s, Parkinson’s, and other dementias, has increased and is expected to double by 2040 (Hickman et al., 2016; Reeve et al., 2014). Anti-aging interventions offer promise in delaying age-related impairments and neurological disease development. The most established anti-aging intervention is calorie-restriction (CR), where caloric intake is reduced by 10-40%. CR consistently increases lifespan and improves health span across model organisms including invertebrates (Lee et al., 2006), rodents (Turturro et al., 1999), and primates (Colman et al., 2014).

In the CNS, CR increases expression of DNA repair and anti-stress proteins (Kisby et al., 2010), improves glucose metabolism efficiency (Willette et al., 2012a), and delays onset of inflammatory cytokines (Swindell, 2009). CR delays neurodegeneration and synaptic dysfunction (Dietrich et al., 2010; Schafer et al., 2015), improves neuroendocrine function (Redman and Ravussin, 2009; Willette et al., 2012b), and promotes induction of genes active in neuroprotection, neural growth (Park et al., 2013), and synaptic and neuronal function (Adams et al., 2008; Wood et al., 2015). CR is also beneficial in alleviating pathology of age-related neurodegenerative diseases, improving cognitive function (Halagappa et al., 2007) and decreasing hippocampal beta-amyloid accumulation in Alzheimer’s models (Wang et al., 2005). Despite these protective effects on brain aging and age-related disease, the molecular mechanisms underlying brain aging processes and those responsive to CR remain elusive.

Epigenetic modifications in the forms of DNA methylation and histone modifications are proposed as central regulators of the aging process (Lopez-Otin et al., 2013), and CR has been proposed to work through epigenetic mechanisms but data supporting this hypothesis is quite limited (Li et al., 2011). DNA methylation is a malleable epigenetic modification in which a methyl group is added to the 5-position of a cytosine ring and plays a complex role in genomic regulation. Proper control of DNA methylation is instrumental to neural differentiation, neural growth, synaptic function and intact cognitive function (Feng et al., 2010; Grigorenko et al., 2016; Lister et al., 2013), key processes that decline with age. Altered methylation at specific genomic locations is associated with the normal aging process (Wyss-Coray, 2016), and recent studies have shown a protective epigenetic effect of anti-aging interventions in the liver and kidney (Cole et al., 2017; Hahn et al., 2017; Kim et al., 2016). We previously described extensive age-related CG and non-CG methylation changes in the hippocampus of old mice (Masser et al., 2017), demonstrating that the DNA modification response to aging is different in the CNS as compared to other organs. However, whether CR affects methylation in the CNS in a pattern consistent with the CNS-specific age-related changes or in a manner like that observed in other tissues is unexplored.

In the current study we investigated whether life-long CR can could prevent DNA methylation changes in the CNS, focusing on the hippocampus, a tissue demonstrating molecular, cellular, and functional impairment with age (Fan et al., 2017). Using a genome-wide sequencing approach, we examined the response of age-related CG and for the first time CH (non-CG) methylation to CR. Additionally, we characterize unique CR-specific changes in both CG and CH methylation, highlighting potential functional similarities and dissimilarities between CG and CH methylation. Lastly, we show that enzymes regulating DNA methylation and demethylation are suppressed by CR but not aging, in part through CR-induced altered promoter methylation, indicating a potential mechanism for CR modulation of the neuroepigenome.

## 2. Materials and methods

### 2.1 Animals

Male C57BL/6 mice were obtained from the NIA aging and caloric restriction colony at 3 and 24 months of age. Mice were housed at the University of Oklahoma Health Sciences Center animal facility and maintained under SPF conditions and individually housed in a HEPA barrier environment. Mice were fed irradiated NIH-31 mouse/rat diet (Teklad, Envigo). CR was initiated at 14 weeks of age at 10% restriction, increased to 25% restriction at 15 weeks, and to 40% restriction at 16 weeks of age. Mice were euthanized by decapitation hippocampi were harvested, snap frozen, and stored at −80°C. Young *ad libitum* (Y-AL, 3 months of age), old *ad libitum* (O-AL 24 months of age) and old calorically restricted (OCR, 24 months of age) samples were collected (n=8/group). All animal experiments were performed according to protocols approved by the OUHSC Institutional Animal Care and Use Committee. Libraries from young and old *ad libitum* animals were used in a previous study and were re-sequenced (Masser et al., 2017).

### 2.2 Library preparation and sequencing

Hippocampal gDNA was isolated by spin columns (Zymo duet) and genome-wide bisulfite capture sequencing in accordance to previously described and validated methods (Masser et al., 2016; 2017). gDNA was quantified by Qubit (Invitrogen) and sheared by sonication (Covaris e220) to an average 200bp fragment size. Fragment size was confirmed by capillary electrophoresis (DNA1000, Agilent). Bisulfite sequencing library preparation was performed with 3µg DNA using SureSelect Mouse Methylseq XT per manufacturer instructions (Agilent). In brief, sheared DNA (n=4/group) was end-repaired, adenylated and adapter ligated prior to bisulfite conversion. Libraries were sized and quantified by capillary electrophoresis to ensure recovery of at least 350 ng of library. Libraries were hybridized to mouse SureSelect Methyl-Seq capture library. Hybridized libraries were captured with Dynabeads MyOne Streptavidin T1 magnetic beads and then bisulfite converted (EZ DNA Methylation-Gold, Zymo Research), amplified according to manufacturer recommendations and purified with AMPure XP beads. Following bisulfite conversion libraries were indexed and confirmed for fragment size by capillary chip (DNA high sensitivity, Agilent). Prior to sequencing, libraries were quantified by PCR (KAPA library quantification kit, Kapa Biosystems), diluted to 4nM, and pooled in equimolar concentrations. Pooled libraries were diluted to 12 pM and then sequenced at 100 bp paired end reads on the Illumina HiSeq2500 platform. Raw fastq files are available through the Sequencing Read Archive (bioproject ID PRJNA397575).

### 2.3 Informatics

Paired-end reads were trimmed using CLC Genomics Workbench v10.1.1. Reads with a Q-score <30 or with >1 ambiguous nucleotide were removed. Reads passing filter were adapter trimmed and an additional 3 bp were removed from the 3’ and 5’ ends of each read. All computational analyses were performed in UNIX and R using custom scripts or published tools (where specified). Trimmed PE reads were aligned as PE to the mouse GRCm38/mm10 assembly (https://genome.ucsc.edu/) using Bismark Bisulfite Mapper version 0.14.4 (Krueger and Andrews, 2011). Following read mapping, methylation calling was performed with Bismark methylation extractor and % methylation was determined by calculating (total C methylated/total C covered) per cytosine in the genome. Only cytosines covered in all groups with a cumulative coverage of 35 were used for analysis, exceeding the coverage recommendations in the field for BS-seq studies (Roadmap Epigenomics et al., 2015; Ziller et al., 2015). Greater than 27M CG and non-CG sites were carried forward for downstream analysis. Differential methylation between groups was determined using Kruskal-Wallis H test in R, significant sites were tested for pairwise multiple comparisons using Conover post hoc test using ‘lsmeans’, and p-values were adjusted for false discovery using the Benjamini-Hochberg procedure. Differentially methylated sites were sites that met statistical cutoff of fdr adjusted p-value < 0.05 and absolute methylation difference > 5%. Age-changes were defined as all differences found between young-AL and old-AL animals (Supplemental Table 1). Diet-changes were defined as all differentially methylated cytosines between AL and CR animals (Supplemental Table 2).

### 2.3 Annotation and enrichment

Coordinates for genic features including exons, introns, and promoters (defined as ±1kb from TSS) were downloaded from UCSC genome browser for GRCm38/mm10 reference genome. CG island shores were defined as 2kb upstream and downstream from the CG island borders and shelves were defined as 2kb upstream and downstream from shores. Coordinates for annotated gene regulatory regions for mouse brain were downloaded from ENSEMBL open database (Zerbino et al., 2015). Finally, coordinates for active histone marks, including H3K27ac, H3K27me3, H3K36me3, H3K4me1, H3K4me3, H3K9me3 and H4K20me1, in the adult mouse hippocampus were obtained from previously published work (Gjoneska et al., 2015). Over- and under-representation of differentially methylated cytosines was determined by overlapping all analyzed sites with the coordinates of genic features, histone marks and gene regulatory regions using ‘bedtools’ (Quinlan and Hall, 2010). Statistical significance of over- and under- representation was determined using hypergeometric test in R. For pathway analysis, genes were annotated for differentially methylated cytosines. The relationship between gene length and number of differentially methylated cytosines was examined in order to eliminate any confounding effects of gene length. No relationship between gene length and number of differentially methylated sites was evident (Supplemental Fig. 1). To control for number of sites analyzed per gene, the ratio of dmCs to total sites analyzed was calculated and genes in the bottom 10^th^ percentile were removed. Analysis of pathways enriched for genes affected for differential methylation was performed with the ‘ReactomePA’ R package. Pathways with broad (>300 genes) or narrow (<7 genes) definitions were excluded post hoc. Additionally, pathways with similar gene lists were consolidated. Complete unedited pathways lists can be found in supplemental tables 3 and 4.

### 2.4 Real-Time PCR (qPCR)

Expression of DNMTs (pan-DNMT1, Mm01151063_m1; E1-DNMT1, Mm00599763_m1; Alt. E1-DNMT1, Mm01174085_m1; DNMT3a1, Mm00432870_m1) and TETs (TET1, Mm01169087_m1; TET2, Mm00524395_m1; TET3, Mm00805756_m1) was determined by real-time qPCR (n=7-8/group) with fluorogenic primer/probes (Life Technologies) as described previously (Mangold et al., 2017a). β-actin (Mm02619580_g1) was used as the endogenous control and each sample was analyzed in technical triplicates.

### 2.4 Experimental Design and Statistical Analysis

Genome-wide methylation analysis was performed with n=4/group. Gene expression experiments were performed with n=7-8/group. All statistical analyses were performed in R. Box plots represent the 25^th^ and 75^th^ percentiles. Differential methylation and enrichment was determined as detailed above. Global methylation differences were compared using linear regression and are written as mean ± SD. Analysis of gene expression using RT-qPCR was compared using One-Way ANOVA with t-test pairwise post-hoc comparisons and Benjamini-Hochberg multiple testing corrections.

## 3. Results

### 3.1 Age related CG methylation is attenuated by life-long caloric restriction

To examine the effect of life-long caloric-restriction on CG methylation in the old brain, hippocampal DNA was isolated from young (3-month-old, Y-AL) and old (24-month-old, O-AL) male mice that were fed *ad libitum* (AL) and old animals fed a 40% caloric restriction (CR) diet (24-month-old, O-CR) starting at 3 months of age. Using genome-wide bisulfite sequencing, base-specific methylation levels across ~2 million CGs were compared between the 3 groups. Global DNA methylation, i.e. the methylation averages across all analyzed CG sites did not change with age or diet (Y-AL: 42.3% ± 0.7; O-AL: 42.6% ± 0.5; O-CR: 42.6%±0.6; p=0.45 by linear regression). Age-associated dmCGs (age-dmCGs) were defined as any methylation difference between young-AL and old-AL animals that met statistical criteria (methylation difference >|5%| and false discovery adjusted p-value < 0.05, for more details see differential methylation in Methods). Comparing young-AL animals to old-AL animals, 41,585 age-related age-dmCGs were identified. Examination of the methylation profile of all age-dmCGs for differences between young-AL and old-CR mice reveals a pattern of regression towards that of young-AL, indicating a ‘younger’ methylation profile in old mice fed a CR diet as compared age-matched *ad libitum* fed mice (Fig. 1A, Supplementary Fig. 2). At single base resolution, life-long CR prevented 32% hypermethylated and 36% hypomethylated age-dmCGs (Fig. 1B). Age-dmCGs showed a higher number of hypermethylated dmCGs than hypomethylated dmCGs. CR-prevented age-dmCGs were proportional to that observed for age-dmCGs (Fig. 1C). Principal component analysis of age-dmCGs unaffected by CR demonstrated clustering of all old mice as compared to young-AL (Fig. 1D). Parallel analysis of CR-prevented age-dmCGs separates young-AL and old-CR from old-AL on the first component, while young-AL and old-CR separate on the second component (Fig. 1E).

**Figure 1.**
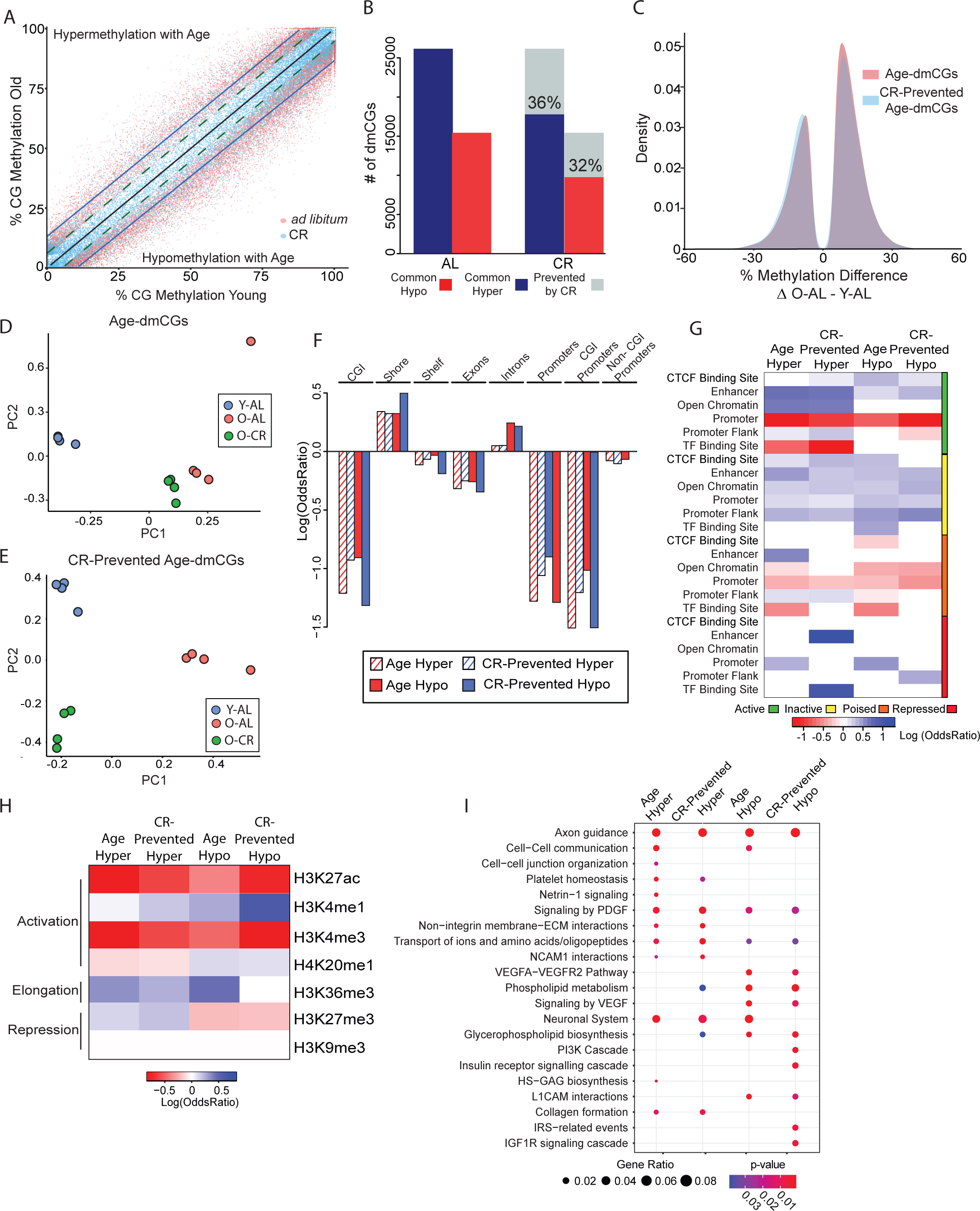
Caloric-restriction prevents age-associated differential CG methylation. **(A)** Scatterplot depicting significant age-dmCG (n = 41,585, adjusted p-value < 0.05 & methylation difference between Old-AL and Young-AL > |5%|) methylation differences between young *ad libitum* (Y-AL) and old *ad libitum* (O-AL) groups (red dots) and young *ad libitum* (Y-AL) and old calorie-restricted (O-CR) groups (blue dots). Linear regression lines pass through Y-AL ~ Y-AL (black), O-AL ~ Y-AL (blue) and O-CR ~ Y-AL (dotted green) age-dmCG methylation values separated by hyper- and hypo-age-dmCGs. **(B)** Quantification of the number of age-dmCGs by hypermethylation (blue) and hypomethylation (red) in animals fed *ad libitum* (AL) or calorie-restricted (CR). Grey indicates age-dmCGs prevented by caloric-restriction (n=13,950). **(C)** Density plot of age-dmCGs (red) and prevented age-dmCGs (blue). PCA plots of mice by age-dmCGs (**D**) and prevented age-dmCGs (**E**). **(F)** Enrichment of age-dmCGs and prevented age-dmCGs in gene-regulatory regions. Red and blue indicate significant under-representation or over-representation (p < 0.05, hypergeometric test) respectively, while white shows no significance (p > 0.05). **(G)** Enrichment of age-dmCGs and prevented age-dmCGs in gene-centric regions, only significant enrichment is shown (p < 0.05, hypergeometric test). **(H)** Enrichment of age-dmCGs and prevented age-dmCGs in annotated histone marks from the adult mouse hippocampus. Red and blue indicate significant under-representation or over-representation (p < 0.05, hypergeometric test) respectively, while white shows no significance (p > 0.05). **(I)** Pathway analysis of genes containing significant age-dmCGs or prevented age-dmCGs. Top enrichment pathways are shown (FDR adjusted p-values < 0.05). Circle size indicates the ratio of genes containing age-dmCGs or prevented age-dmCGs in a pathway, color indicates p-values.

The localization of differentially methylated sites by genomic context and their interaction with other epigenomic marks can aid in understanding the role of DNA methylation changes in genomic regulation. Age-dmCGs were mapped to known coordinates of gene-centric features to annotate the enrichment of differentially methylated sites. Similar patterns of enrichment are evident across genic and CG island (CGI) features for hyper- and hypo-methylated age-dmCGs and age-dmCGs prevented by CR (Fig. 1F). Age-dmCGs and CR-prevented age-dmCGs were enriched in introns and CGI shores while under-represented in islands, CGI shelves, exons, and promoters. Depletion of age-dmCGs in promoters was more pronounced in promoters containing CGIs. Previous studies have reported that DNA methylation may alter gene expression through interaction with gene regulatory regions (Roadmap Epigenomics et al., 2015). To identify enrichment at specific gene regulatory regions dmCGs were compared to the ENSEMBL brain specific regulatory build (Zerbino et al., 2015) which was constructed using sequencing methods to identify CTCF binding sites, enhancers, open chromatin regions, promoters, promoter flanks, and transcription factor binding sites and their specific activation states (active, inactive, poised, repressed). Enrichment of age-dmCGs varied by type of regulatory region and activation state (Fig. 1G). Notable was enrichment in all inactive state gene regulatory regions. Inactive regions are defined as genomic regions not associated with CTCF, DNase hypersensitivity, or ChIP-Seq chromatin peaks from the ENSEMBL brain database, and therefore indicate that differential methylation at these regions is independent of interaction with other known epigenetic marks in the brain. Age-dmCGs were also enriched in active state regulatory regions and occurred in CTCF binding sites (hypo only), enhancers (hypo and hyper), open chromatin regions (hyper only), and promoter flank (hyper only). CR prevented age-dmCGs generally demonstrated a similar enrichment pattern. Using previously published data characterizing histone modification in the adult mouse hippocampus (Gjoneska et al., 2015) we identified significant enrichment of age-dmCGs and CR prevented sites associated with both repressive and activational histone marks (Figure 1H). The strongest under-representation was in H3K27ac and H3K4me3 regions associated with activated genes, while the strongest over-representation was in activational H3K4me1 and repressive H3K36me3 marks.

The interaction between gene expression and DNA methylation is still an active area of exploration with the canonical view of an inverse relationship between DNA methylation and gene expression has been challenged both in neurons (Sharma et al., 2016) and following CR (Hahn et al., 2017). Therefore, potential effects of DNA methylation on gene expression were made without assumptions as to whether methylation is positively or negatively regulating a specific gene. Hypermethylated age-dmCGs were found in 2,060 genes and hypomethylated age-dmCGs were found in 1,622. Of those genes, CR prevented hypomethylated age-dmCGs in 745 genes and hypermethylated age-dmCGs in 1,028 genes. Pathways involved in neural cell communication, neural cell growth, cellular integrity, cell metabolism and inflammatory pathways were affected by both hyper- and hypomethylation with aging (Fig. 1H, supplemental table 3). Pathways enriched for prevented age-dmCGs by CR were mostly regulatory pathways involving cell metabolisms such as phospholipids metabolisms and IGF signaling as well as pathways regulating neuronal cellular structure and integrity. These findings are consistent with the negative effect of aging on neuronal integrity and positive cellular effect of CR on neuronal survival and growth (Hornsby et al., 2016; Seib and Martin-Villalba, 2015).

### 3.2 Age related CH methylation (CC, CA, and CT) is attenuated by life-long caloric restriction

The effect of CR on CH methylation has not been previously examined. CH methylation is higher in the brain relative to other tissues (Lister et al., 2013), is differentially regulated as compared to CG methylation (He and Ecker, 2015; Masser et al., 2017), and has unique functional impacts on gene regulation in the brain (Lister and Mukamel, 2015). We analyzed >25M CHs sites and tested if age-dmCHs can also be prevented by CR. Absolute methylation level of most dmCHs was generally lower than that of CGs. Average global CH methylation levels did not change with age or with diet (Y-AL: 1.59% ± 0.07; O-AL: 1.64% ±0.05; O-CR: 1.58%±0.8; p=0.83 by linear regression). More age-dmCHs (79,056) were identified compared to age-dmCGs. The methylation profile of age-dmCHs in old animals fed CR diet was closer to that of young-AL animals compared to that of old animals fed *ad libitum* (Fig. 2A). The number of hypermethylated age-dmCHs was higher than hypomethylated dmCHs in a manner similar to CGs. Life-long CR prevented 59% of all hypermethylated age-dmCHs and 11% of hypomethylated age-dmCHs (Fig 2B). Unlike CGs, prevention of hypermethylated age-dmCHs by CR was 10 times more likely than CR prevention of hypomethylated age-dmCHs (Fig. 2C). Principal component analysis of age-dmCHs unaffected by CR shows a distinct clustering of both AL and CR old animals compared to young-AL animals (Fig. 2D). Clustering of samples by the methylation profile of CR-prevented age-dmCHs shows that old-AL animals separate from young-AL and old-CR animals (Fig. 2E).

**Figure 2.**
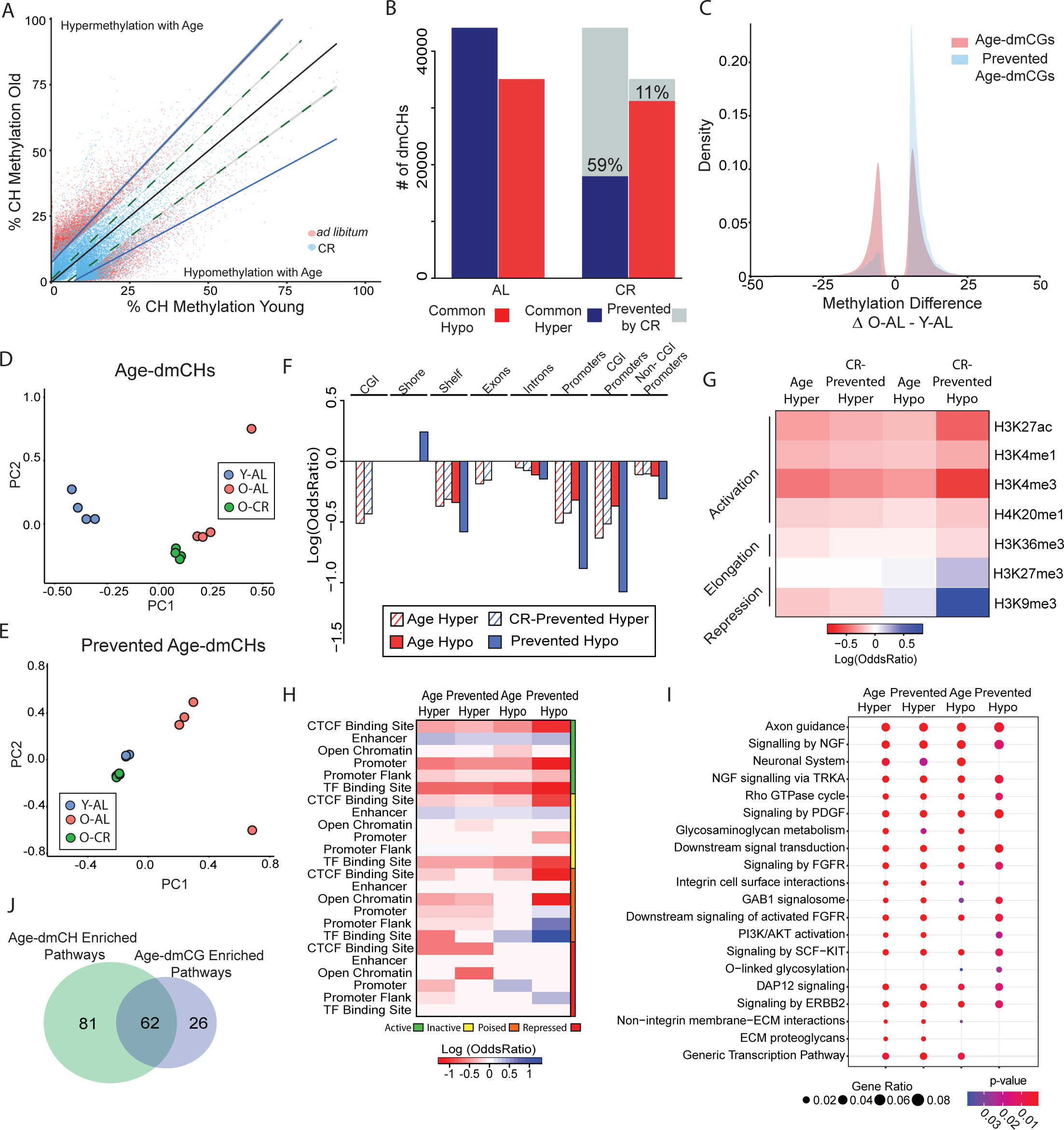
Caloric-restriction prevents age-associated differential CH methylation. **(A)** Scatterplot depicting significant age-dmCH (n = 79,056, adjusted p < 0.05 & methylation difference between Old-AL and Young-AL > |5%|) methylation differences between young *ad libitum* (Y-AL) and old *ad libitum* (O-AL) groups (red dots) and young *ad lib* (Y-AL) and old calorie-restricted (O-CR) groups (blue dots). Linear regression lines pass through Y-AL ~ Y-AL (black), O-AL ~ Y-AL (blue) and O-CR ~ Y-AL (dotted green) age-dmCH methylation values separated by hyper- and hypo- age-dmCHs. **(B)** Quantification of the number of age-dmCHs by hypermethylation (blue) and hypomethylation (red) in animals fed *ad libitum* (AL) or calorie-restricted (CR). Grey indicates age-dmCHs prevented by caloric-restriction (n = 29,880). **(C)** Distribution plot of age-dmCHs (red) and prevented age-dmCHs (blue). PCA plots of samples by age-dmCHs (**D**) and prevented age-dmCHs (**E**). **(F)** Enrichment of age-dmCHs and prevented age-dmCHs in gene-centric regions, only significant enrichments are shown (p < 0.05, hypergeometric test). **(G)** Enrichment of age-dmCHs and prevented age-dmCHs in annotated histone marks from the adult mouse hippocampus. Red and blue indicate significant under-representation or over-representation (p < 0.05, hypergeometric test) respectively, while white shows no significance (p > 0.05). **(H)** Enrichment of age-dmCHs and prevented age-dmCHs in gene-regulatory regions. Red and blue indicate significant under-representation or over-representation (p < 0.05, hypergeometric test) respectively, while white shows no significance (p > 0.05). **(I)** Pathway analysis of genes containing significant age-dmCHs or prevented age-dmCHs. Top enrichment pathways are shown (FDR adjusted p-values < 0.05). Circle size indicates the ratio of genes containing >2 age-dmCHs or prevented age-dmCHs in a pathway, color indicates p-values. **(J)** Overlap between pathways enriched for age-dmCHs and age-dmCGs.

Differentially methylated CHs were generally under-represented in CG islands and shelves, and in genecentric elements. The exception being an enrichment of prevented hypomethylated age-dmCH in CGI shores (Fig. 2F). Taken together, under-representation of dmCHs in CG islands and genic regions indicate that changes in methylation with age, and those prevented by CR, occur primarily in intragenic regions and are partially excluded from CG rich regions. Age-dmCHs were generally not associated with specific histone peaks (log odds ratio <0). The only observed enrichment was of hypomethylated age-dmCHs in H3K9me3, a pattern quite dissimilar from age-dmCGs (Fig. 2G).

Enrichment analysis of age-dmCHs in gene-regulatory regions reveals that age-dmCHs and prevented age-dmCHs, regardless of gain or loss in methylation, are most highly enriched in both active and inactive enhancers (Fig 2H). Hypomethylated age-dmCHs and CR-prevented age-dmCHs were also enriched in poised TF binding sites and repressed promoters. Pathways enriched for genes affected by age-dmCH were functionally similar to those enriched by age-dmCG, including pathways involved in metabolic regulation, cellular signaling, and regulation of neural cell structure and growth. CR-prevented age-dmCHs were enriched in pathways primarily involved in energetics, metabolism and neurite growth (Fig. 2I, Supplementary Table 4). Of the pathways found to be affected by differential methylation, 62 were common to age-dmCHs and age-dmCGs (Fig 2J).

### 3.3 CR induces chances in methylation at sites unaffected by age

Caloric-restriction was sufficient to prevent >1/3 of all age-dmCs (CGs and CHs). CR also induced DNA methylation changes independent of aging as evident when comparing old AL mice and to old CR mice. Stated differently, the majority of sites where methylation is regulated with CR do not have a change in methylation with aging on AL feeding (Fig. 3A). Of all differentially methylated dmCGs between AL and CR, 69% were induced by CR independent of an age effect (Fig. 3B), as exemplified in Supplementary Figure 3A. More CR-specific hypermethylation effects (CR-dmCGs) than hypomethylated sites were observed (Supplementary Fig 3B).

**Figure 3.**
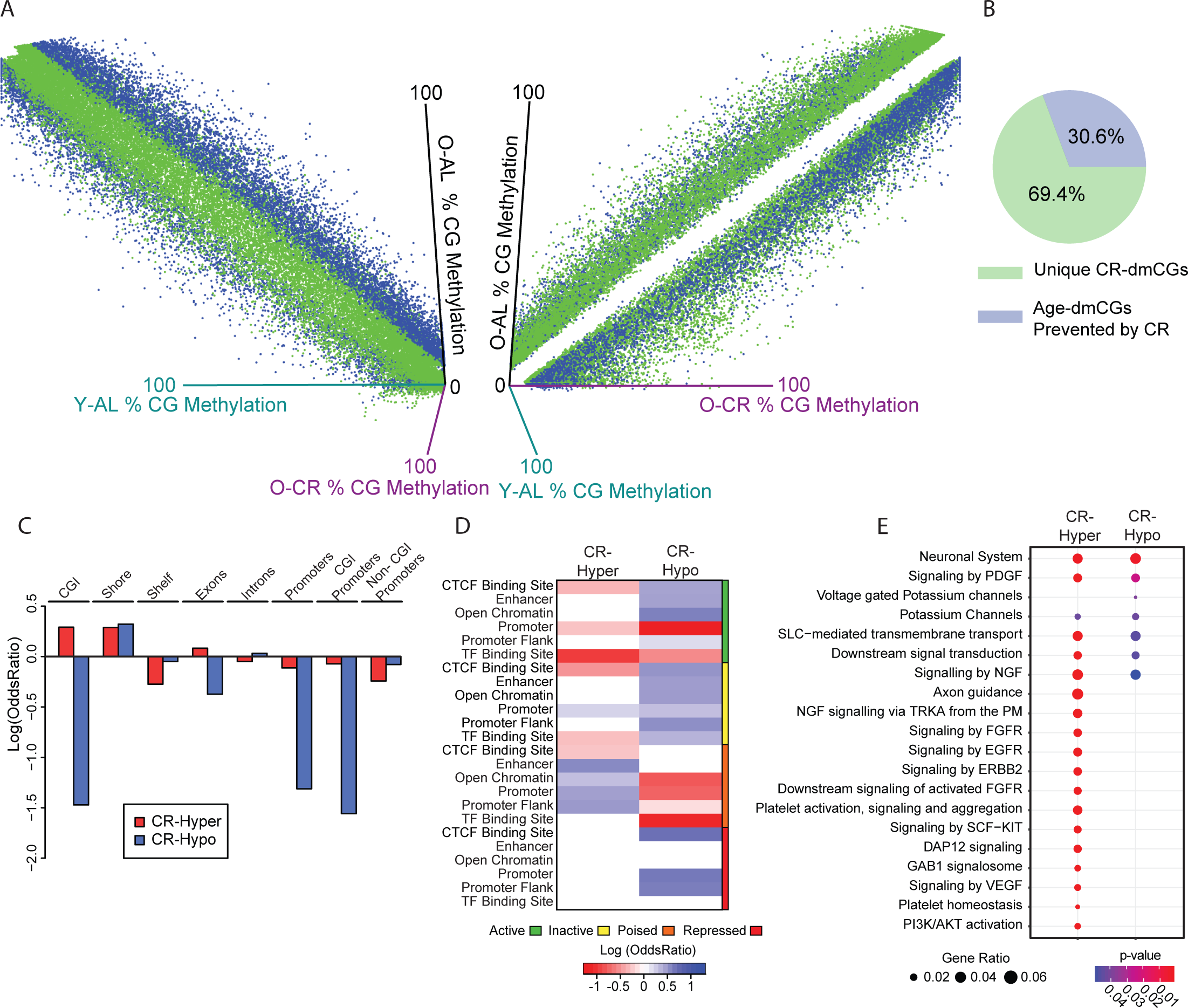
CR induces changes in CG methylation independent of aging. **(A)** *Left*: 3D scatterplot showing the methylation levels of significant CR-dmCGs including unique CR-dmCGs (green) and age-dmCGs prevented by CR (blue) (adjusted p < 0.05 & methylation difference between Old-AL and Old-CR > |5%|) focusing on young-AL and old-AL. *Right*: 90° rotation on the Y-axis of the 3D plot showing that unique CR-dmCGs (green) regress away from the midline when comparing old-AL and Old-CR methylation levels. **(B)** Proportion of diet-dmCGs unique to CR (green) and those with an age interaction (prevented age-dmCGs, blue) (N = 46,813). **(C)** Enrichment of CR-dmCGs in gene-centric regions (p < 0.05, hypergeometric test). **(D)** Enrichment of CR-dmCGs in gene-regulatory regions. Red and blue indicate significant under-representation or over-representation (p < 0.05, hypergeometric test) respectively, while white shows no significance (p > 0.05). **(E)** Pathway analysis of genes containing >2 significant diet-dmCGs. Top enrichment pathways are shown (FDR adjusted p-values < 0.05). Circle size indicates the ratio of genes containing diet-dmCGs in a pathway, color indicates p-values.

The genomic localization of CR-dmCGs is markedly different from age-dmCGs. Hypermethylated CR-dmCGs were over-represented in CG islands and shores and under-represented in CG shelves, while hypomethylated CR-dmCGs were only enriched in shores (Fig. 3C). With respect to genic regions CR-dmCGs were primarily under-represented with the exception of over-representation of hypermethylated sites in exons and hypomethylated sites in introns. The same distinction between hyper- and hypo-methylated CR-dmCGs was observed in gene regulatory regions. Enrichment of CR-dmCGs was primarily in poised regulatory regions while enrichment of hypomethylated CR-dmCGs was observed in active regulatory features, including CTCF binding sites, enhancers, open chromatin and promoter flanks (Fig. 3D). Active promoters were not enriched for hyper- and hypo-methylated CR-dmCGs; however, hypermethylated CR-dmCGs were enriched in inactive and repressed promoters. Consistent with the enrichment patterns of CR-dmCGs in regulatory regions, hypermethylated CR-dmCGs were enriched in repressive H3K9me3 and H3K27me3 marks (Zhang et al., 2015), while hypomethylated CR-dmCGs were over-represented in both activational and repressive marks (Supplementary Fig 3C). Genes affected by CR-dmCGs were enriched for pathways involving energy regulation, inflammatory pathways, senescence-associated secretory phenotype, and phagocytosis (Fig. 3E, Supplementary Table 3).

### 3.4 CR induced changes in CH sites are clustered

The CR-specific effect on methylation was also observed in non-CG sites (Fig. 4A). Changes in CH methylation with CR occurred generally at lower methylation levels (< 50%, Fig. 4A – right panel) and did not change with age (Fig. 4A, B). Twice as many hypermethylated (64,191 sites) than hypomethylated CR induced-dmCHs (26,237 sites) were observed, similar to CR-dmCGs. The average magnitude of CR induced methylation was higher in hypermethylated CH sites than hypomethylated CH sites (10% vs −8%, Fig. 4C).

**Figure 4.**
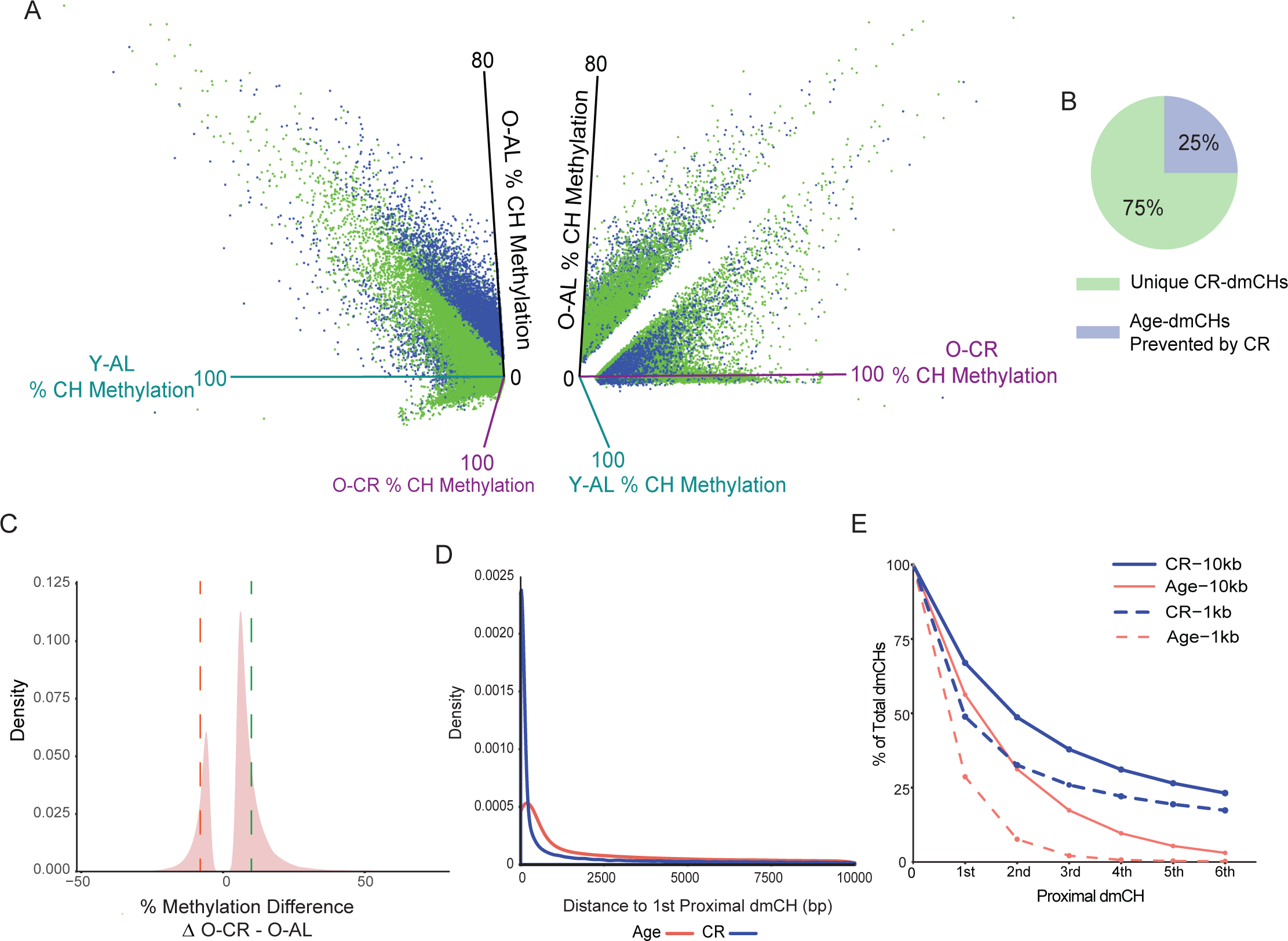
CR induced differential CH methylation is clustered at specific genomic loci. **(A)** *Left*: 3D scatterplot showing the methylation levels of significant CR-dmCHs including unique CR-dmCHs (green) and age-dmCHs prevented by CR (blue) (adjusted p < 0.05 & methylation difference between Old-AL and Old-CR > |5%|) focusing on young-AL and old-AL. *Right*: 90° rotation on the Y-axis of the 3D plot showing unique CR-dmCH (green) regress away from the midline when comparing old-AL and Old-CR methylation levels. **(B)** Proportion of CR-dmCHs unique to CR (green) and those with age interaction (prevented age-dmCHs, blue) (N = 90,428). **(C)** Distribution of CR-dmCHs. Dotted lines mark the average methylation differences of hypomethylated CR-dmCHs (red, 8%) and hypermethylated CR-dmCHs (green, 10%). **(D)** Distribution plot comparing the proximity between dmCHs by age and CR focusing on 10 kb segments. **(E)** Line graph representing % of dmCHs by degree of proximity within 10kb (solid lines) or 1kb (dotted line) by age or by CR.

CG dinucleotides density is highest in CG islands and promoter regions and lower in other genomic regions. On the other hand, the distribution of CH dinucleotides is relatively uniform across the genome (He and Ecker, 2015). Since CHs tend to be sporadically distributed across the genome, clustering of CH differential methylation with age or with CR may be indicative of their upstream regulation. Computing the distance between dmCHs with age or CR revealed that the majority (62%) of all dmCHs (age and CR) occurred within 10kb of another dmCH. Focusing on 10kb segments, CR-dmCHs were in closer proximity to each other than age-dmCHs (Fig. 4D, E). The closer proximity to the next dmCH is more pronounced difference between CR-dmCHs than age-dmCHs as evident within 1kb windows (Figure 4E, dashed lines).

### 3.5 CR induced-dmCHs are over-represented in CGI-promoters and depleted from non-CGIs promoters

Hypermethylation of CR-dmCHs was over-represented in CG islands, shores and exons, while hypomethylation was over-represented in introns, a localization pattern similar to CR-dmCGs (Fig. 5A). Hypermethylated diet-dmCHs were also over-represented in promoters containing CGIs but not in non-CGI promoters. Using the ENSEMBL regulatory build, we further divided promoters by their activation state and found that hypermethylated CR-dmCHs are enriched in active and poised promoters and promoter flanks, and are under-represented in repressed and inactive promoters (Fig. 5B). Enrichment of hypomethylated CR-dmCHs was mainly observed in inactive gene regulatory elements. CR-induced hypomethylation was under-represented all histone marks while CR-hypermethylation was associated with both repressive and active chromatin marks (Supplementary Fig. 4A). This is consistent with the enrichment of CR induced CH hypermethylation in promoters and active promoters as well as supporting the possibility that changes in CH methylation induced by CR might result in altered gene transcription.

**Figure 5.**
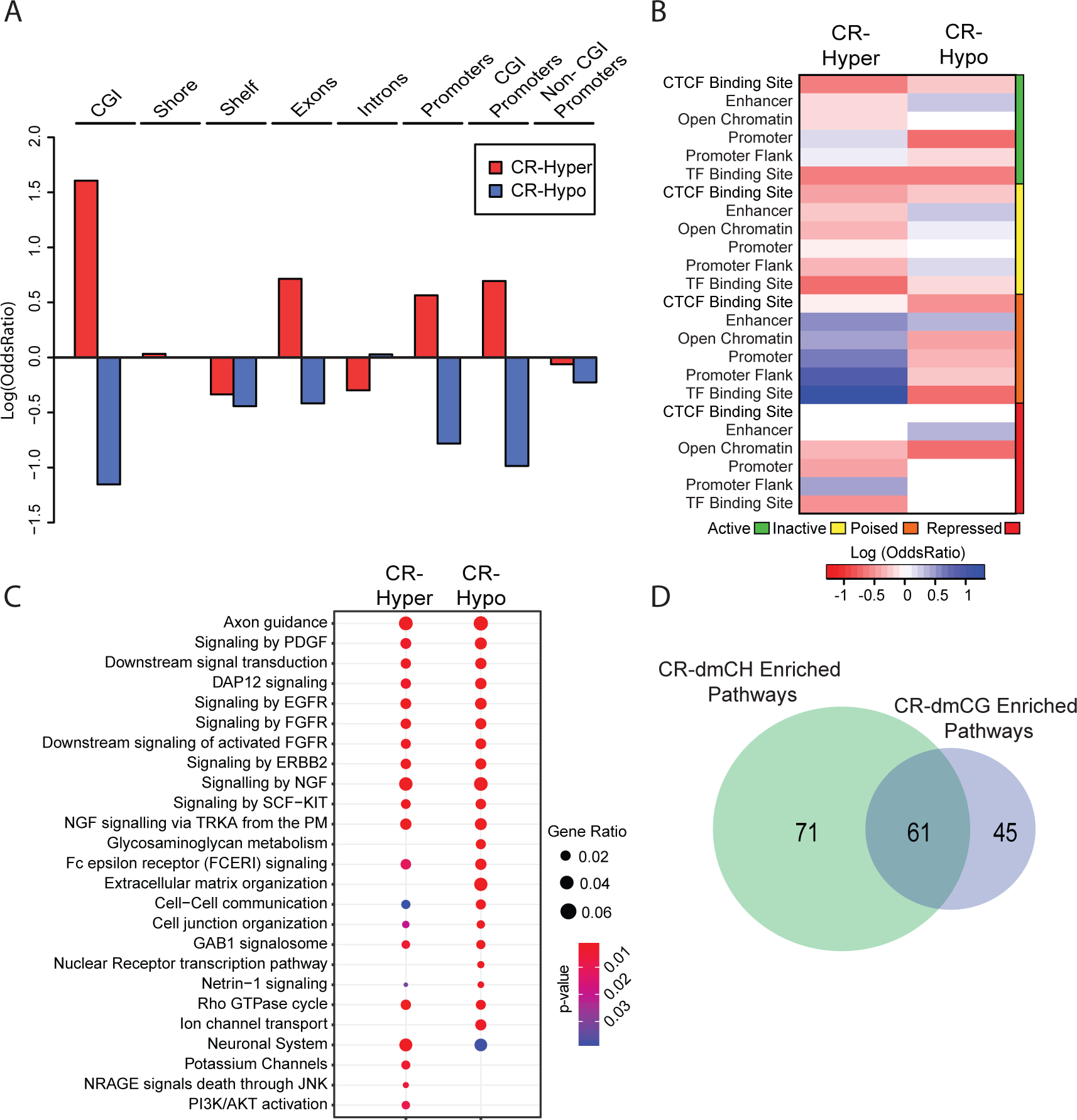
CR-induced differential CH methylation enrichment patterns. **(A)** Enrichment of CR-dmCHs in gene-centric regions, only significant enrichment is shown (p < 0.05, hypergeometric test). **(B)** Enrichment of CR-dmCHs in gene-regulatory regions. Red and blue indicate significant under-representation or over-representation (p < 0.05, hypergeometric test) respectively, while white shows no significance (p > 0.05). **(C)** Pathway analysis of genes containing >2 significant CR-dmCHs. Top enrichment pathways are shown (FDR adjusted p-values < 0.05). Circle size indicates the ratio of genes in pathway containing CR-dmCHs, color indicates p-values. **(D)** Overlap between pathways enriched for CR-dmCHs and CR-dmCGs.

Higher numbers of genes contained CR-induced CH hypermethylation (2,606 genes) than hypomethylation (1,978 genes), consistent with the distribution of CR-dmCHs. Top enrichment pathways included those regulating cell structure, neural cell communication and growth, and energy metabolism. Inflammatory pathways were mostly enriched for genes containing hypomethylated dmCHs (Fig. 5C, Supplementary Table 4). Of the 177 pathways affected by both CR-dmCHs or CR-dmCGs, 61 were found in common including pathways regulating inflammation and neuronal integrity (Fig. 5D).

### 3.6 CR results in changes in the expression of enzymes regulating DNA methylation

CR-dmCGs were enriched in the DNA methylation pathway (Supplemental Table 3), and mapped to DNMT and TET promoters. Specifically, hypermethylated CR-dmCs, 2 dmCGs and 4 dmCHs in TET3, and 3 dmCGs and 6 dmCHs in DNMT1 were observed (Fig. 6A-B). A commensurate CR-specific decrease in TET3 mRNA level was evident (Fig. 6C). For DNMT1 no change in expression was evident with a pan-isoform assay, but when isoform-specific expression corresponding to an alternate the start sites around the CR-dmCs were examined, an isoform-specific decrease in expression was observed (Fig 6D). CR-specific decreases in DNMT3a1 and TET2 mRNA expression were observed while TET1 was unchanged with aging or CR (Fig 6F).

**Figure 6.**
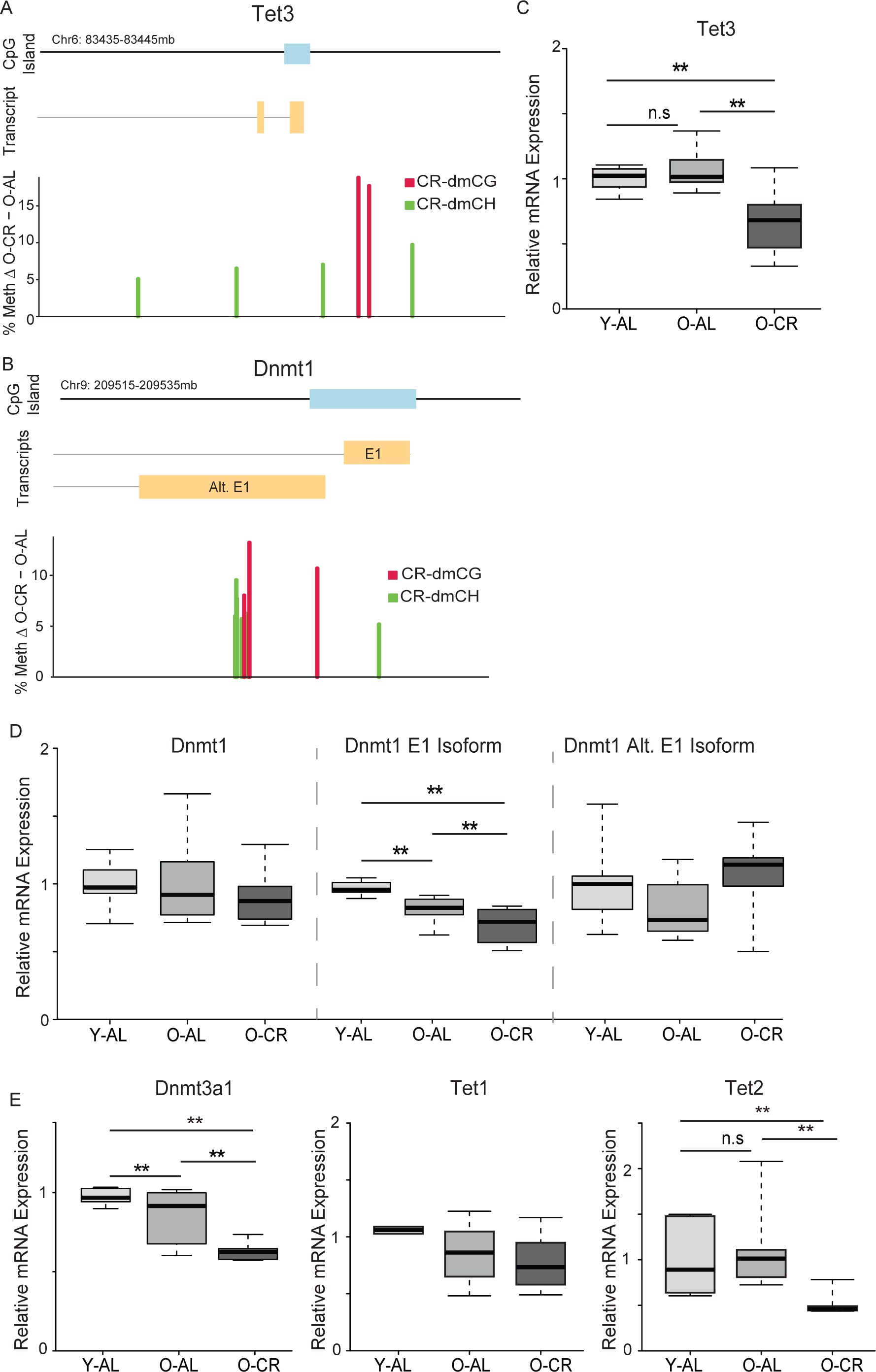
CR-induced differential methylation of DNA regulatory enzymes. Calorie-restriction induces hypermethylation of CGs (red) and CHs (green) in the promoter regions of TET3 **(A)** and DNMT1 **(B)**. **(C)** mRNA expression of TET3 decreases with CR but not age. **(D)** mRNA expression of DNMT1 using a pan-isoform assay does not change with CR or aging, however, when isoform-specific expression is analyzed the exon 1 (E1) isoform decreases with age and diet while the isoform using the alternative start site (Alt. E1) is unchanged with age or CR. **(E)** mRNA expression of DNMT3a1 decreases with diet and age. TET1 expression did not change in any condition and TET2 expression decreased with diet but not age. n=7-8/group, *p < 0.05, **p < 0.001, One-way ANOVA with Benjamini-Hochberg correction).

## 4. Discussion

The beneficial neurological effects of CR are well-characterized in both aging and age-related neurodegenerations (Van Cauwenberghe et al., 2016). This study examined, for the first time, whether CR could prevent age-related changes in CNS DNA methylation, consistent with the concept that methylation is a potential molecular mechanism by which CR slows brain aging. CR has been reported outside of the CNS (liver, kidney, and blood) to prevent age-related CG methylation changes (Cole et al., 2017; Hahn et al., 2017; Kim et al., 2016; Maegawa et al., 2017) but has not been examined in the CNS. We have reported that the predominant form of methylation changes with age in the CNS are in non-CpG (CH) contexts (Masser et al., 2017) and the effects of CR on CH methylation have not been described in any tissue. Therefore, we examined whether CR could prevent age-related differential CG and CH methylation. Our findings demonstrate that prevention of age-related differential methylation by CR is observed in the brain, in both CG and CH contexts, and with enrichment in specific genomic elements and genes related to fundamental aging processes. We also identified CR-specific effects on CG and CH methylation patterns that may contribute to the neuroprotective effects of CR. By including CH methylation in our analysis, we provide the novel finding that CR attenuation of age-related DNA methylation changes in not exclusive to CG methylation but also occurs in CH context, an observation that may be unique to neural tissues. These CH findings stand in contrast to the patterns of age-related changes and their prevention by CR observed outside the CNS further demonstrating the unique nature of the neuroepigenomic response to aging

### 4.1 Prevention of Aging-Associated Methylation Patterns

DNA methylation changes in the old hippocampus are observed in both CG and CH contexts (Masser et al., 2017). Over a third of all age-related differentially methylated cytosines in either context were prevented by CR. The distribution of prevented age-related CG differential methylation was proportional to that of age-dmCGs. However, in CH dinucleotides, CR preferentially prevented age-related hypermethylation. This preferential prevention of hypermethylation may be caused by the CR-specific reduction in DNMT3a1 expression observed. DNMT3a is required for *de novo* methylation of CH sites (Guo et al., 2014) and of the two DNMT3a isoforms, DNMT3a1 and not DNMT3a2 is expressed in the mature hippocampus (Hadad et al., 2016). Expression of DNMTs and TETs is responsive to CR in the liver (Hahn et al., 2017) suggesting that this could be conserved response to CR across tissues. However, no other data has been reported on CH methylation in response to aging or CR in other tissues. As non-CpG is required for transcriptional regulation of genes involved in synaptic and neural growth (Chen et al., 2015; Jang et al., 2017), prevention of age-related CH changes could be critical to the neuroprotective effects of CR.

Changes in dmCG and dmCH methylation with aging, and those prevented by CR, demonstrated distinct patterns across genic and CpG island regions, chromatin, enhancers, and other genomic elements. Several salient points emerged from these findings: 1) dmCGs and dmCHs prevented by CR demonstrate distinct patterns of over- and under-representation from each other, 2) CR-prevented dmCGs and dmCHs follow the enrichment of age-related changes specific to the type of site, and 3) patterns of CG enrichment are dissimilar from those reported outside the CNS. On this later point, preferential prevention of age-related CG hypomethylation (Cole et al., 2017) has been reported in the liver, while here we observed an equivalent proportion of age-related CG hyper- and hypo-methylation prevented by CR. In the blood, promoter CpG islands are hypermethylated and non-promoter CpG islands are hypomethylated (Maegawa et al., 2017) with aging. This is typical of the proposed changes in methylation with aging (Booth and Brunet, 2016) but are not observed here in the CNS with aging or prevention of age-related changes by CR. The functional implications remain to be determined but it is possible to speculate that the CNS, which is predominantly composed of long-lived post-mitotic cells, is markedly different from other tissues like liver and blood which are composed of mitotic and comparatively short-lived cells that are being replenished throughout the lifespan. This does not obviate the possibility of specific differentially methylated sites with aging that are conserved between tissues (Horvath, 2013). Our findings also demonstrate that prevention of age-related changes by CR in CG methylation are reproducibly observed across mouse models and tissues.

While there are no other datasets of age-dmCHs prevented by CR to which our findings can be compared, the patterns of prevention (and also thereby aging) are significantly different from those observed in CG contexts. This raises important questions of how changes in methylation are directed to specific loci in a base-specific manner. The regulatory mechanisms, such as DNMT and TET co-factors, that direct changes to specific genomic locations is only beginning to be understood (Song et al., 2015; Wu and Zhang, 2017) and is generally unexplored in the CNS. Targeting of DNMTs and TETs to specific genomic loci can be directed through post translational histone modifications at specific genomic locations (Balasubramanian et al., 2012; Cedar and Bergman, 2009; Lehnertz et al., 2003) and may be altered by CR, thus changing recruitment of DNMTs and TETs. While we did not characterize changes in histone modification, DNA methylation changes with both age and CR were co-localized with various histone marks previously mapped in the adult hippocampus. In particular, regions demonstrating loss of methylation with aging were associated with H3K4me1 (Fernandez et al., 2015), a histone mark indicating poised enhancers (Calo and Wysocka, 2013). Also of note was that the highest enrichment of prevented CG hypomethylation changes at H3K9me3 sites. Prevention of H3K9 methylation has been associated with improved brain function with aging (Snigdha et al., 2016). DNA methylation and histone modifications symbiotically regulate each other (Cedar and Bergman, 2009) and in future studies combined data from chromatin and methylation from aged subjects is needed to further investigate the interplay between the two.

Age-related dmCs occurred in promoters and intragenic regions of genes enriched in inflammatory, neuronal integrity, neuronal function, and synaptic transmission pathways similar to those previously identified as altered with aging (Keleshian et al., 2013; Mangold et al., 2017a; 2017b; Meng et al., 2016; Stilling et al., 2014) and CR (Zeier et al., 2011) in gene expression studies. Similar pathways were enriched for methylation changes prevented by CR, indicative of the neuroprotective actions of CR in the CNS. Transcriptional regulation by DNA methylation is an ongoing area of investigation. Studies examining the interactions between gene expression and differential methylation with age and CR (Hahn et al., 2017) and in the brain (Sharma et al., 2016) suggest a far more complex relationship than the simple inverse correlation generally assumed. While demonstrating causality between differential methylation of thousands of sites and gene expression remains unfeasible we hypothesize that methylation changes with age contribute to the dysregulation of genes in these pathways. How changes with age or prevention by CR in regions such as open chromatin, CGI shores and CTCF binding sites contribute to regulating gene expression during aging require greater understanding of the three dimensional organization of the genome. Further investigation is required to discover the many intricate mechanisms by which methylation acts as a gene expression regulator with aging – studies preferably performed on isolated cellular populations as patterns of methylation vary by cell type in the CNS (Lister et al., 2013).

### 4.2 CR-Specific Changes in Methylation

The majority of CR-induced differentially methylated sites were independent of aging. The preferential CG hypermethylation of CR-specific dmCGs is shared by findings in the liver (Hahn et al., 2017). These CR-specific changes demonstrated different distributions in promoters, genes, and enhancers compared to age-related dmCs prevented by CR. For example, CR-specific effects could serve to counter balance over-/under-activation of a gene or pathway with aging that is driven by non-epigenetic mechanisms. Striking was the finding that CR-specific changes in CH methylation are not sporadic throughout the genome but are clustered at higher density than age-associated changes. CR-induced gain of methylation at CH loci was most evident in regions where CH methylation is rare, such as promoters, CG islands, TF binding sites and active enhancers (Guo et al., 2014; Laurent et al., 2010; Lister et al., 2009; Xie et al., 2012). The regulation and function of this clustering is not readily apparent. Methylation of CHs appears to respond differently than CGs and likely serves a non-redundant role in genomic regulation because 1) enrichment of dmCGs and dmCHs was different at promoters, TF binding sites, enhancers and open chromatin and 2) the affinity of the DNMTs and TETs to CHs is lower than to CGs (He and Ecker, 2015; Ramsahoye et al., 2000; Suetake et al., 2003). The differential localization of dmCHs as compared to dmCGs provides evidence that different, yet unknown, targeting mechanisms are being used to regulate CH and CG methylation with diet.

Several pathways are known to be upregulated by food-restriction and its mimetics (Fusco and Pani, 2013). Prior gene expression studies in the brain have shown that mTOR signaling and insulin signaling are downregulated by CR, as they are in other tissues, while brain-specific induction of pathways including calcium signaling, synaptic vesicle regulation, neurotrophic signaling, neural growth, differentiation and function, and neuronal integrity are also observed (Schafer et al., 2015; Wood et al., 2015). We observed CR-induced methylation changes in pathways previously reported to change following CR in the brain, specifically calcium signaling, energy metabolism, NGF signaling, FGFR signaling, and pathways involved in axon guidance and growth. While a direct effect was not established, the common findings between our study and previous gene expression studies suggest a role for DNA methylation as a possible mechanism for the neuroprotective effect of CR against brain aging. This is exemplified by enrichment of diet-dmCGs in DNA methylation and senescence-associated secretory phenotype pathways; both pathways have been proposed as potential regulators of the aging process (Lopez-Otin et al., 2013). Indeed, we demonstrate decreased expression of DNMT1 and TET3 accompanies hypermethylation of these genes.

These findings provide a rationale for future studies to address several important questions raised. Discovery and examination of DNMT and TET targeting co-factors are needed to demonstrate mechanisms by which changes are directed to specific genomic regions. Studies of the time course of differential methylation with aging as well as the duration and age of CR initiation are needed in both males and females as this study only examined males. As age-related methylation differences demonstrate different patterns between males and females it will be of interest to see whether there are sex-specific preventative effects. Additionally, inclusion of hydroxymethylation analysis through bisulfite sequencing will provide further and more specific insight into how DNA modifications change with aging and may be prevented or reversed with anti-aging treatments. These questions combined with the need to examine specific cellular populations and to examine specific subregions of the hippocampus which we and others have demonstrated to have specific responses to aging and CR (Masser et al., 2014; Zeier et al., 2011).

### 4.3 Conclusions

Taken together our data supports a role of calorie-restriction in maintaining cellular homeostasis through preventing changes in DNA methylation with aging. Changes in methylation were enriched in pathways shown to improve following CR and that are beneficial to brain aging. A significant CR-specific response was evident in addition to simply preventing age-related changes that may also serve a neuroprotective role, including autoregulation of DNMTs and TETs. These findings also demonstrate the extensive modulation of non-CpG/CH methylation by CR. Compared to other tissues the response of CNS methylation to aging and CR is markedly different and indicative of tissue-specific epigenetic responses to CR.

## Acknowledgments

The authors would like to acknowledge Laura Blanco-Berdugo for computational advice and the University of Oklahoma Supercomputing Center for Education & Research (OSCER) for allocating computational resources used for data analysis. This work was supported by the Donald W. Reynolds Foundation, the Oklahoma Center for Neuroscience translational seed grant, the Oklahoma Nathan Shock Center of Excellence in the Biology of Aging Targeted DNA Methylation and Mitochondrial Heteroplasmy Core (P30AG050911), the National Institute on Aging (R01AG045693, T32AG052363), and National Eye Institute (R21EY024520, R01EY021716).

---

**Supplementary Figure 1.** Analysis of the interaction between differential methylation to gene length. The ratio between dmCs to sites analyzed is poorly correlated (Pearson’s r < 0.1) to gene length in both CG (**A**) and CH (**B**) context.

**Supplementary Figure 2.** Representatives of an age-dmCG (left) and a prevented age-dmCG (right).

**Supplementary Figure 3 (A)** Single base example of a unique CR-dmCG. **(B)** Distribution of CR-dmCGs. Dotted lines mark the average methylation differences of hypomethylated CR-dmCGs (red, −12.5%) and hypermethylated CR-dmCGs (green, 12.3%). **(C)** Enrichment of CR-dmCGs in annotated histone marks from the adult mouse hippocampus. Red and blue indicate significant under-representation or over-representation (p < 0.05, hypergeometric test) respectively, while white shows no significance (p > 0.05).

**Supplementary Figure 4** Enrichment of CR-dmCHs in annotated histone marks from the adult mouse hippocampus. Red and blue indicate significant under-representation or over-representation (p < 0.05, hypergeometric test) respectively, while white shows no significance (p > 0.05).

## Notes

**Conflict of interest:** The authors declare no competing financial interests

